# Thalamic inputs determine functionally distinct gamma bands in mouse primary visual cortex

**DOI:** 10.1101/2020.07.09.194811

**Authors:** Nicolò Meneghetti, Chiara Cerri, Elena Tantillo, Eleonora Vannini, Matteo Caleo, Alberto Mazzoni

**Affiliations:** The Biorobotics Institute, Scuola Superiore Sant’Anna, 56025 Pisa, Italy; Department of Excellence for Robotics and AI, Scuola Superiore Sant’Anna, 56025 Pisa, Italy; Neuroscience Institute, National Research Council (CNR), via G. Moruzzi 1, 56124 Pisa, Italy; Fondazione Umberto Veronesi, Piazza Velasca 5, 20122 Milan, Italy; Department of Pharmacy, University of Pisa, Via Bonanno Pisano 6, 56126 Pisa, Italy; Fondazione Pisana per la Scienza Onlus (FPS), via Ferruccio Giovannini 13, San Giuliano Terme, 56017 Pisa, Italy; Scuola Normale Superiore, Piazza dei Cavalieri 7, 56100 Pisa, Italy; Department of Biomedical Sciences, University of Padua, viale G. Colombo 3, 35131 Padua, Italy

## Abstract

Gamma band is known to be involved in the encoding of visual features in the primary visual cortex (V1). Recent results in rodents V1 highlighted the presence, within a broad gamma band (BB) increasing with contrast, of a narrow gamma band (NB) peaking at ∼60 Hz suppressed by contrast and enhanced by luminance. However, the processing of visual information by the two channels still lacks a proper characterization. Here, by combining experimental analysis and modeling, we prove that the two bands are sensitive to specific thalamic inputs associated with complementary contrast ranges. We recorded local field potentials from V1 of awake mice during the presentation of gratings and observed that NB power progressively decreased from low to intermediate levels of contrast. Conversely, BB power was insensitive to low levels of contrast but it progressively increased going from intermediate to high levels of contrast. Moreover, BB response was stronger immediately after contrast reversal, while the opposite held for NB. All the aforementioned dynamics were accurately reproduced by a recurrent excitatory-inhibitory leaky integrate-and-fire network, mimicking layer IV of mouse V1, provided that the sustained and periodic component of the thalamic input were modulated over complementary contrast ranges. These results shed new light on the origin and function of the two V1 gamma bands. In addition, here we propose a simple and effective model of response to visual contrast that might help in reconstructing network dysfunction underlying pathological alterations of visual information processing.

**Significance Statement:** Gamma band is a ubiquitous hallmark of cortical processing of sensory stimuli. Experimental evidence shows that in the mouse visual cortex two types of gamma activity are differentially modulated by contrast: a narrow band (NB), that seems to be rodent specific, and a standard broad band (BB), observed also in other animal models.

We found that narrow band correlates and broad band anticorrelates with visual contrast in two complementary contrast ranges (low and high respectively). Moreover, BB displayed an earlier response than NB. A thalamocortical spiking neuron network model reproduced the aforementioned results, suggesting they might be due to the presence of two complementary but distinct components of the thalamic input into visual cortical circuitry.

## Introduction

Gamma band ([30-100] Hz) synchronization is a widespread functional mode in the mammalian cortex, known to improve long-range information transmission (Konig et al., 1995; Sohal, 2016). Coherently, gamma synchronization contributes to the processing of sensory information, from audition (Brosch et al., 2002; Tsunada and Eliades, 2020) to olfaction (Beshel et al., 2007; Lepousez and Lledo, 2013), touch (Siegle et al., 2014) and nociception (Tan et al., 2019; Heid et al., 2020).

The role of gamma synchronization in the processing of visual stimuli has been investigated for decades (Gray and Singer, 1989; Gray et al., 1989). Indeed, visual stimulus features are well known for modulating the power and the central frequency of neocortical oscillations in the gamma range. Previous studies adopting different animal models (mice, cats, human and non-human primates) have highlighted gamma power to be critically dependent on orientation (Gray and Singer, 1989; Berens, 2008; Onorato et al., 2020), size (Gieselmann and Thiele, 2008; Perry et al., 2013), speed (Gray et al., 1989; Friedman-Hill, 2000; Womelsdorf et al., 2006), direction (Liu and Newsome, 2006), and contrast (Logothetis et al., 2001; Henrie and Shapley, 2005; Ray and Maunsell, 2010; Saleem et al., 2017; McAfee et al., 2018; Bartoli et al., 2019).

Cortical gamma band oscillations originate from the coordinated interaction of excitation and inhibition (Sohal et al., 2009) and might be detected by Local Field Potential (LFP), even though single unit activity is irregular (Buzsáki and Wang, 2012). Many modeling works have described how gamma synchronization originates in excitatory-inhibitory networks and how it is affected by external stimuli (Wilson and Cowan, 1972; Leung, 1982; Ermentrout and Kopell, 1998; Brunel and Wang, 2003; Börgers and Kopell, 2003; Geisler et al., 2005). Later, several later studies focused on how such networks could capture contrast-induced modulation of gamma band in primary visual cortex (Mazzoni et al., 2008, 2010, 2011; Battaglia and Hansel, 2011; Jia et al., 2013; Roberts et al., 2013; Barbieri et al., 2014; Lowet et al., 2015; Zachariou et al., 2019). These works captured a variety of properties of gamma band in the visual cortex of primates, but recent evidence showed that, at least in mice, the gamma band might be composed of two types of resonances with specific frequency ranges and different encoding properties. A very narrow gamma band close to 60 Hz has been observed in the visual cortex of awake mice (Niell and Stryker, 2010; Lee et al., 2014). In a recent work (Saleem et al., 2017) the broad band activity in the 30–90 Hz range was found to co-exist with this narrow band oscillation at 60 Hz. Crucially, while the broad band power increased with contrast, as in primates, the narrow band power increased with luminance and decreased with contrast (Saleem et al., 2017). Neurons highly synchronized at 60 Hz were observed in the Lateral Geniculate Nucleus (Saleem et al., 2017) and the retina (Storchi et al., 2017), suggesting a subcortical origin for the cortical narrow band. Recent modeling works reproducing in detail the structure of the mouse visual cortex (Arkhipov et al., 2018; Billeh et al., 2020) did not specifically address this issue. Overall, the function of the mouse gamma narrow band in the primary visual cortex and the mechanisms underlying its modulation still need to be properly clarified.

To address this issue, we stimulated awake mice with gratings of different contrasts and performed a spectral analysis of the resulting V1 LFP, showing that narrow and broad gamma band not only have opposite but even complementary sensitivity range. Simulations with recurrent spiking network accurately reproduced the behavior of both narrow and broad gamma band and support the hypothesis that the origin of the former might be due to a periodic component embedded within the thalamic input.

## Materials and Methods

### Experimental design

#### Mice

Experiments were conducted in accordance with the European Community Directive 2010/63/EU and were approved by the Italian Ministry of Health. Animals were housed in a 12 h light/dark cycle with food and water available *ad libitum*. Adult (4–6 weeks old) C57BL/6J female mice were used in all experiments.

#### Recording implant

Animals were chronically implanted with a custom-made aluminum head post, and a rectangular recording chamber (2 x 1.5 mm) of dental cement (Ivo-clar Vivadent Inc., USA) was built over the primary visual cortex (i.e. between 0 and 1.5 mm anterior and between 1.5 and 3.5 lateral to the lambda suture) leaving the skull intact. A ground electrode was placed over the cerebellum. The electrode was connected to a pin socket and secured to the skull by acrylic dental cement. Surgery was conducted under deep avertin anaesthesia (7 ml/kg; 20% solution in saline, i.p.; Sigma Aldrich). Animals were then allowed to recover for 3 days. Following recovery, animals were habituated for 3 days to the head fixation apparatus. A craniotomy overlying primary visual cortex was performed 24 hours before the first recording session. To preserve the cortical surface, the recording chamber was filled with a layer of agar (Sigma Aldrich, USA) and the silicone elastomer Kwik (World Precision Instrument, USA.) as a protective cap.

#### Extracellular recordings in awake mice

Recordings were performed on awake mice. Mice were carefully placed in the head fixation apparatus. After removing the protective cap, the recording chamber was filled with sterile saline solution (0.9%) to preserve and moisten the tissue.

A NeuroNexus Technologies 16-channel silicon probe (**Figure 1 A**) with a single-shank (A1×16-3mm-50-177) was mounted on a three-axis motorized micromanipulator and slowly lowered into the visual cortex (in the central region of the recording chamber) till the depth of 1000 µm. Before the beginning of the recording, the electrode was allowed to settle for about 5 min. The neurophysiological data were continuously recorded using a 16-channel Omniplex system (Plexon, Dallas, TX). At the end of the extracellular recording session, the recording chamber was covered with the protective cap as described above. Each animal underwent two recording sessions on two different days.

**Figure 1.**
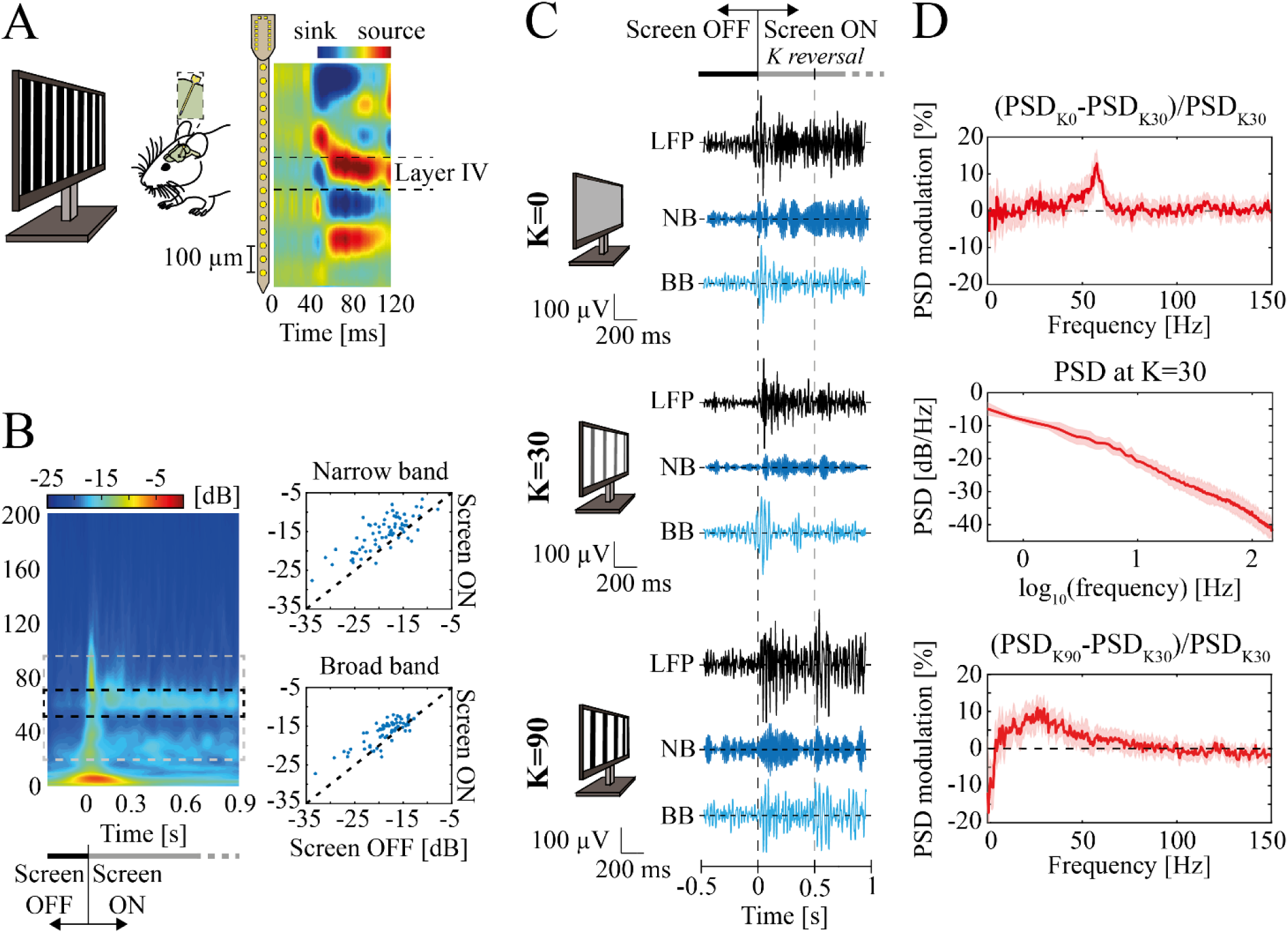
Experimental setup and data. ***A***, (*left*) Representative scheme of the experimental design. Square-wave 1 Hz alternating gratings at different contrast levels were used for visual stimulation. A linear 16-channels probe (with 50 μm spacing between electrodes) was inserted into the mouse primary visual cortex. (*right*) Mean across animals of CSDs aligned by the earliest current sink. ***B***, (*left*) Mean scalogram for K=0 within -200 ms to 900 ms around screen onset. The dashed rectangle depicts the frequency bands ranges: narrow band (middle, black) and broad band (black). (*right*) Scalogram magnitude comparison between screen OFF (from -200 to 0 ms) and screen ON condition (from 0 to 900 ms) in high NB (top) and BB (bottom). Statistical differences were accounted for by the Wilcoxon’s matched pairs signed rank test. For both gamma bands, p-values were far less than 0.001. ***C***, Examples of filtered local field potential recorded in mice V1 while viewing a uniform gray screen (top), or alternating gratings at contrast K=30 (middle) and K=90 (bottom). The examples are reported between [- 0.5; 1] s around screen onset. Examples were band-passed between (i) 10 to 100 Hz (black traces) just for representative purposes; (ii) 45 to 65 Hz to display the NB; (iii) from 20 to 45 Hz and from 65 to 90 Hz to display the BB range. Dashed lines indicate screen onset (black) and the first contrast reversal (grey). Monitors’ sketches schematically represent visual contrast. ***D***, LFP modulation of minimal contrast (i.e., K=0) (top) and maximal contrast (i.e., K=90) (bottom) with respect to the power spectral density (PSD) at K=30 (middle). Modulation is defined as the difference between the power of a frequency at a given contrast level (K=0 or K=90 in this case) with the power at reference contrast K=30, normalized to the latter power. Shaded regions indicate S.E.M.

All the electrophysiological data were processed using custom-written MATLAB codes (The MathWorks, Natick, MA, USA). The extracellular signals were sampled at 1 kHz and band-pass filtered from 1 to 500 Hz in pre-processing.

#### Visual stimuli

Visual stimuli were computer-generated using the Matlab Psychophysics Toolbox with gamma correction and presented on a display (Sony; 40 9 30 cm; mean luminance 15 cd/m^2^) placed 25 cm from the head of the mouse (**Figure 1 A**), covering the center of the visual field. Extracellular signals were recorded in response to 30 abrupt reversals (1 Hz) of vertical square-wave gratings (spatial frequency, 0.06 c/deg; contrast or K ranging from 0 to 90%). Signals were amplified (5000-fold), bandpass filtered (0.5–500 Hz), and fed into a computer for storage and analysis. For each recording 30 contrast reversal were presented to the head-fixed animals (n=12 for K<50 and n=7 for K=90). Pooling animals together, we performed: 71 recordings at K=0; 61 recordings at K=10 and K=20; 100 recordings at K=30; 66 recordings at K=50; 32 recordings at K=90.

### Neurophysiological data analysis

#### CSD analysis and Layer IV identification

For each channel, visual evoked potential waveforms related to contrast reversals were extracted from the LFPs by signal averaging. For each recording session, current source density (CSD) was computed by applying a standard algorithm (according to the second spatial derivative estimate of the laminar LFP time series, (Haberly and Shepherd, 1973; Freeman and Nicholson, 1975)) along with the iCSD toolbox for MATLAB (Pettersen et al., 2006). A value of 0.3 S/m was taken as a measure of cortical conductivity (Gaussian Filter: standard deviation = 0.05 mm). Layer IV was identified in each recording session (two for each animal) with the channel corresponding to the earliest current sink.

#### Spectral analysis

From extracellular recordings, we extracted Local Field Potentials (LFPs) by low pass filtering at 200 Hz (see **Figure 1 C** for filtered LFP examples). LFPs were z-scored prior to spectral analysis. The power spectral density (PSD) of the z-scored LFPs was computed with the Fast Fourier Transform via the Welch method (pwelch function in Matlab), dividing the time window under investigation into sub-windows of 500 ms with 50% overlap. The only exception was when investigating the temporal structure of narrow and broad band PSD (**Figure 2 C**): z-scored LFPs were segmented in consecutive temporal windows after each contrast reversal. PSDs were, consequently, estimated independently for each of them and averaged across trials. Given the short duration of the time windows (200 and 300 ms) the Welch method was applied with no sub-window. Since the contrast *K* = 30 minimized both narrow and broad band gamma power (see Results), we expressed the spectral response at other contrasts as the spectral modulation relative to the response to this stimulus (**Figure 1 D**, top and bottom):

**Figure 2.**
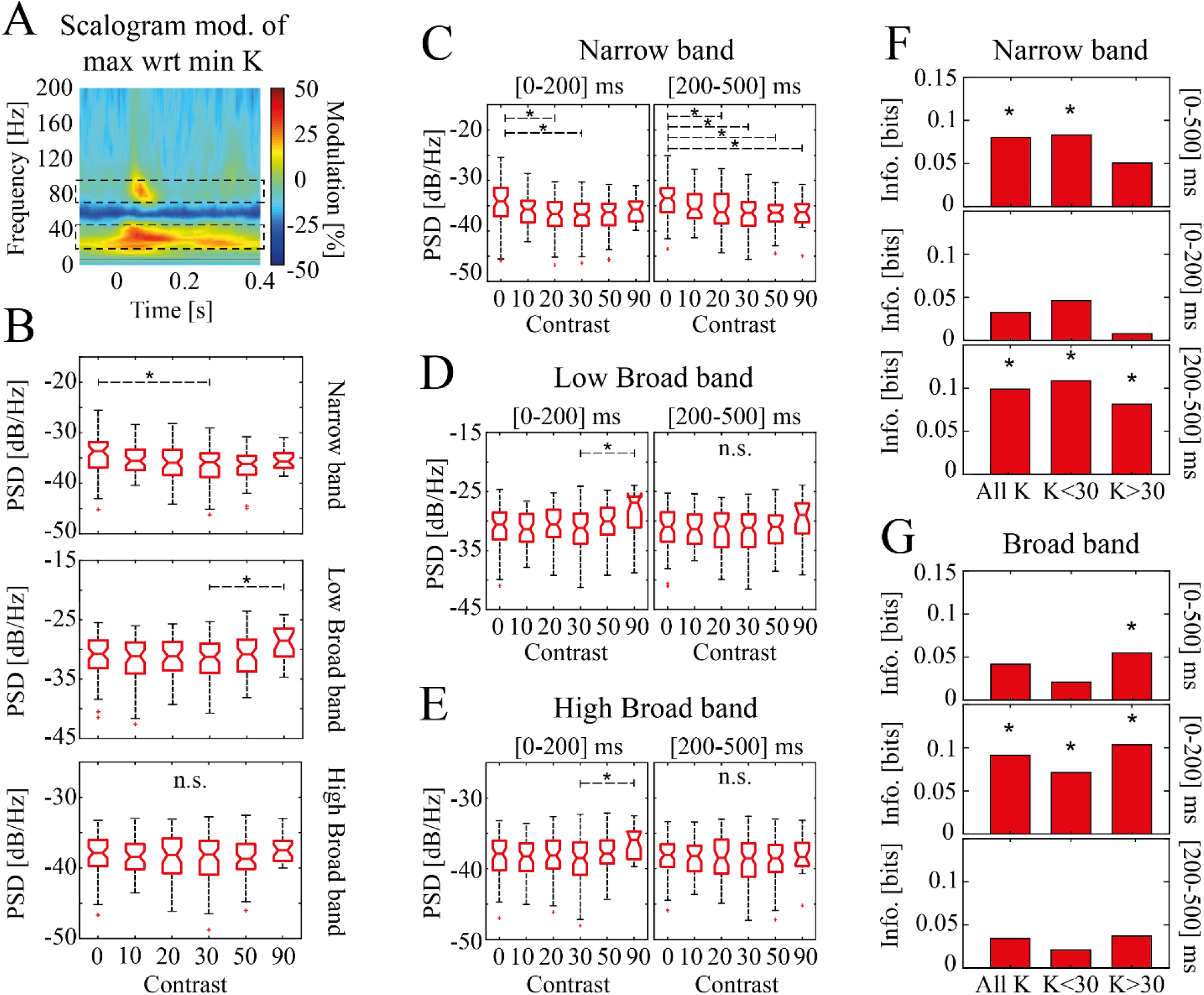
Narrow and broad band gamma contrast-driven modulation in layer IV of mouse V1. ***A***, K=90 vs K=0 modulation of LFP scalogram, pooled across trials of all animals (see Materials and methods). Dashed black rectangles indicate the broad band (BB) composed of two ranges: low BB ([45-65] Hz) and high BB ([65-90] Hz). These two bands are separated by the narrow band (NB, [45-65] Hz). ***B***, PSD of NB (top), low BB (middle), high BB (bottom) as a function of visual contrast pooled across all animals and recordings. On each box, the central mark indicates the median, bottom and top edge indicate the 25th and 75th percentiles. The whiskers extend to the most extreme data points not considered outliers (which are represented by red crosses). Asterisks indicate significant difference. ‘n.s.’ indicates the lack of statistical difference overall possible pairs. ***C***, PSD of NB for [0-200] ms (left) and [200-500] ms (right) after contrast reversal. On each box, the central mark indicates the median across animals, bottom and top edge indicate the 25th and 75th percentiles. The whiskers extend to the most extreme data points not considered outliers (which are represented by pluses). Asterisks indicate significant difference whereas ‘n.s.’ stands for non-significant statistical difference. ***D***, Same as (***C***) for low broad band. ***E***, Same as (***D***) for high broad band. ***F***, Information carried by PSD of narrow band (see Materials and Methods) about all contrast levels (‘All K’), low range of contrasts (‘K<30’), and high range of contrasts (‘K>30’). PSD modulation was considered for the whole inter-contrast-reversal interval ([0-500] ms, top), for 200 ms following contrast reversal ([0-200] ms, middle), or for a time window spanning [200-500] ms after contrast reversal (bottom). Asterisks indicate mutual information values exceeding the significance threshold (p<0.05; bootstrap test). ***G***, Same as (***F***) for broad band.

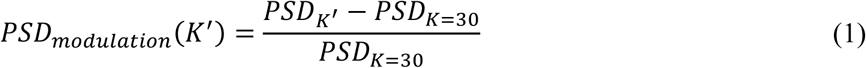

Where *PSD*_*K*=30_ is the median LFP PSD across recordings and animals when contrast K=30 is presented (**Figure 1 D**, middle) and *PSD*_*K*_′ is the median LFP PSD across recordings and animals when contrast K=K’ is presented.

We investigated the evolution in time of the LFP by means of wavelet analysis. LFP scalograms were computed using the continuous wavelet transform (*cwt* function in Matlab), using the analytic Morse wavelet with symmetry parameter equal to 3 and the time-bandwidth product equal to 60. Scalograms were separately computed for each experimental recording and then split into 500 ms long consecutive time windows: specifically, from -100 to 400 ms around every square-wave contrast reversal. We decided not to include in the analysis the LFP evoked by the visual stimulus onset (i.e., the first 500 ms of the recorded LFP after stimulus onset) since it induced flash-like response as the pre-stimulus consisted in a dark screen (**Figure 1 B**). As for the PSD modulation analysis, the mean scalogram of the LFPs at K=30 (**Figure 5 A**) was the reference time-frequency map on relatively to which we computed the modulation of the other contrast levels. The scalogram modulation was computed by averaging the modulated time-frequency maps of every recording at a given contrast level (splitted in 500 ms windows around the grating reversal instants as described above). We computed then the median of the modulated scalogram of every recording within the narrow or the broad band to compute the time evolutions of these two gamma bands.

To define the broad and narrow band frequency limits, we computed the modulation of the mean scalogram across trials and animals of K=90 with respect to K=0 (**Figure 2 A**). We defined broad band (BB) the set of frequencies for which the modulation exceeded the 20%, i.e., the two disjoint ranges [20-45] Hz and [65-95] Hz. We defined narrow band (NB) the gamma band interval between these two ranges, i.e., [45-65] Hz. Next, we evaluated the onset latency of narrow band activation. We computed, for every recording, the onset time of the inter-trial median NB modulation amplitude (Bartoli et al., 2019). First, for each NB signal, we marked the first time point at which the NB amplitude exceeded a threshold (< 25^th^ percentile for at least 40 ms in the time window between -100 to 400 ms; ∼40% of the trials were discarded as they did not meet this criterion). Next, a 200 ms wide window was extracted around that time point (100 ms before and 100 after). This window was segmented into 50 ms bins with 80% overlap and a linear regression was fit to each bin. The first time-point of the bin with the highest slope and smallest residual error was defined as the onset of the narrowband gamma oscillations.

The same procedure was repeated to estimate broad band onset latency, with a threshold of >75^th^ percentile due to the opposite polarity of the modulation.

#### Spectral information analysis

In order to determine how well the PSD modulation of the narrow band encoded the contrast level of visual stimuli, we computed the mutual information (Shannon, 1948) I(K; NB_mod_) carried by the narrow band PSD modulation with respect to K=30 throughout the whole inter-contrast-reversal interval, NB_mod_, about the set of contrasts K:

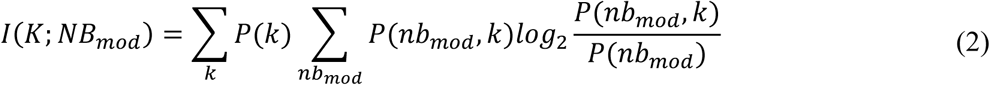

where *P*(*K*) is the probability of contrast K to be presented, *P*(*nb*_*mod*_) is the probability distribution of the narrow band modulation over all the contrasts, and *P*(*nb*_*k*_, *k*) is the probability of the NB modulation *nb*_*mod*_ to be observed when contrast stimulus K is presented. The probabilities, both marginal and conditional, were computed by discretizing the power modulations into 10 equipopulated bins. The same analysis was performed for the broad band. Due to the small number of trials for single animals, probabilities were estimated over all trials and recordings.

Similarly, we also computed the mutual information carried by NB and BB when considering the PSD modulation within 200 ms or from 200 ms to 500 ms following contrast reversal.

We further investigated the amount of information carried by the two gamma bands about low or high contrast levels (low: K<30 and high: K>30). Low contrast information was computed as described above, after random permutation of high contrast-response association (average over 500 permutations), and vice-versa.

We also analyzed the amount of synergistic interaction between narrow and broad band modulations when jointly considered:

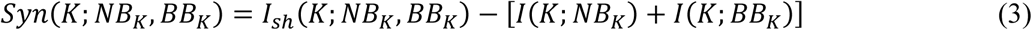

where *I*_*sh*_ is the information computed after randomly permuting, independently for each response, the order of trials collected in response to a stimulus.

Limited dataset bias was accounted for by applying the Panzeri-Treves correction (Panzeri and Treves, 1996). Significant information (p=0.05) was estimated using the 95th percentile of bootstrap information. All information quantities were computed in Matlab with Information Breakdown Toolbox (Magri et al., 2009).

### Network model

#### Spiking network model of mice primary visual cortex

The cortical network structure was the same used in a string of previous works (Mazzoni et al., 2008, 2010, 2011), to which we refer for a full description. Briefly, the simulated network is composed of *N* = 5000 leaky integrate and fire (LIF) neurons (Tuckwell, 1988): 80% excitatory neurons with AMPA-like synapses, and 20% inhibitory neurons with GABA-like synapses (Braitenberg and Schüz, 1991). The network is sparse and random, the connection probability between any directed pair of cells being 0.2 (Sjöström et al., 2001; Holmgren et al., 2003) (**Figure 3 A**, middle).

**Figure 3.**
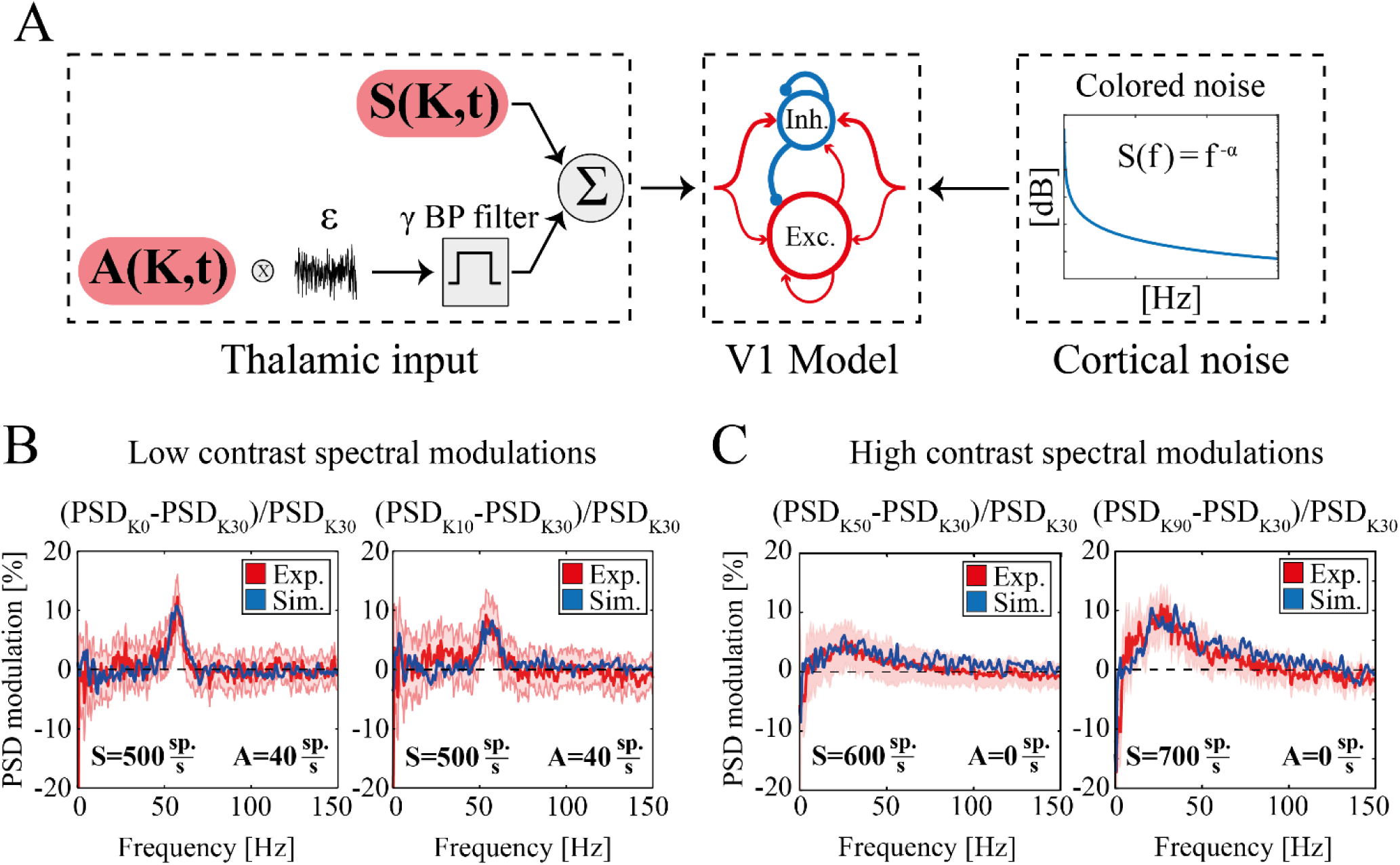
Experimental and simulated spectral modulation at different visual contrasts. ***A***, Schematic representation of the model V1 network. From left to right: (i) thalamic inputs; sustained component S(K,t) (top) and oscillatory component of amplitude A(K,t) (bottom); (ii) sparse leaky integrate and fire network of excitatory (n=4000, red) and inhibitory neurons (n=1000, blue). The size of the arrows represents schematically synaptic strength. In addition to recurrent interactions, both populations receive external excitatory inputs from thalamus and cortex; (iii) colored noise (see Materials and Methods) modeling ongoing unstructured cortical activity. ***B***, Modulation of LFP PSD for K=0 (left) and K=10 (right) with respect to K=30 in experimental (red) and simulated (blue) data. The red shaded area represents SEM of experimental data across animals and recordings. In both simulations sustained thalamic input was set to S=500 sp./s, while the oscillatory input was set to A=40 sp./s for K=0 (left) and A=30 sp./s for K=10, (right). ***C***, Modulation of LFP PSD for K=50 (left) and K=90 (right) with respect to K=30 in experimental (red) and simulated (blue) data. The red shaded area represents SEM of experimental data across animals and recordings. In both simulations, oscillatory thalamic input was set to A=0 sp./s, while the sustained input was S=600 sp./s for K=50 (left) and S=700 sp./s for K=90 (right).

The membrane potential *V*^*k*^ of each neuron *k* evolves according to (Brunel and Wang, 2003):

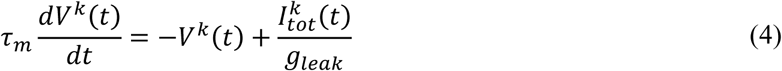

where *τ*_*m*_ is the membrane time constant, *g*_*leak*_ is the leak membrane conductance (see Table 1 for values) and 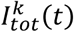 is the total synaptic input current. The latter was given by the sum of all the synaptic inputs entering the k-th neuron:

**Table 1.** Synaptic parameters.

|  | GABA on inhibitory | GABA on excitatory | AMPA<sub>recurrent</sub> on inhibitory | AMPA<sub>recurrent</sub> on excitatory | AMPA<sub>external</sub> on inhibitory | AMPA<sub>external</sub> on excitatory |
| --- | --- | --- | --- | --- | --- | --- |
| $g_{syn}[\text{nS}]$ | 2.700 | 2.010 | 0.233 | 0.178 | 0.317 | 0.234 |
| $\tau_l[\text{ms}]$ | 1 | 1 | 2 | 2 | 2 | 2 |
| $\tau_r[\text{ms}]$ | 1 | 1 | 0.2 | 0.4 | 0.2 | 0.4 |
| $\tau_d[\text{ms}]$ | 5 | 5 | 1.25 | 2.25 | 1.25 | 2.25 |

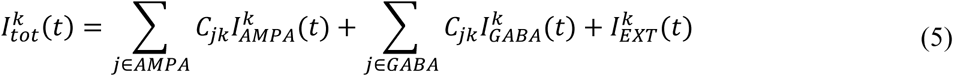

Where *C*_*jk*_ ≠ 0 if j projects to k, and 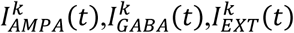 the different synaptic inputs entering the k-th neuron from recurrent AMPA, GABA, and external AMPA synapses respectively. The synaptic inputs currents were modeled as:

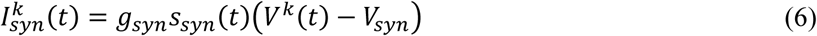

where *g*_*syn*_ are the synaptic conductances (Markram et al., 1997; Gupta, 2000; Bartos et al., 2001, 2002) (see Table 1), and *V*_*syn*_ are the reversal potential of the synapses (V_GABA_=-80 mV and V_AMPA_=0 mV). The function *s*_*syn*_(*t*) described the time course of the synaptic currents and depends on both the synapse type and on the kind of neuron receiving the input. Specifically, every time a pre-synaptic spike occurred at time *t*^*^, *s*_*syn*_(*t*) of the postsynaptic neuron was incremented by an amount described by a delayed difference of exponentials (Brunel and Wang, 2003):

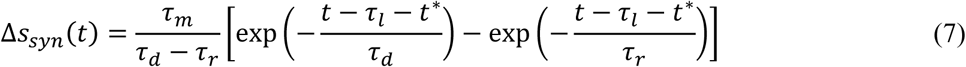

where the latency *τ*_*l*_, the rise time *τ*_*r*_ and the decay time *τ*_*d*_ are listed in Table 1 for each synapse type (Xiang et al., 1998; Zhou and Hablitz, 1998; Angulo et al., 1999; Gupta, 2000; Kraushaar and Jonas, 2000; Bartos et al., 2001).

The LFP of the simulated network was estimated as the sum of the absolute value of the GABA and AMPA currents (both external and recurrent) that enter all excitatory neurons (Mazzoni et al., 2015) as in previous works involving modeling of the primary visual cortex (Mazzoni et al., 2008; Barbieri et al., 2014). Simulated LFP was analyzed with the same procedures followed for experimental LFP (see above).

#### External inputs to the simulated network

For all neurons the external input 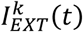 is the sum of two terms: a noisy excitatory external input representing the activity from thalamocortical afferents (**Figure 3 A**, left) and colored noise mimicking stimulus-unspecific cortical activity (Mazzoni et al., 2010) (**Figure 3 A**, right). This simulated external input was implemented as a series of spike times that activated excitatory synapses with the same kinetics as recurrent AMPA synapses, but different strengths (Table 1). These synapses were activated by independent realizations of random Poisson spike trains, with a time-varying rate identical for all neurons. This time-variant rate was given by:

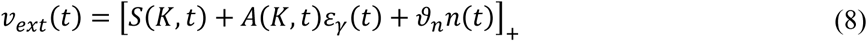

where *K* is the contrast level, *S*(*K, t*) the sustained input rate, *A*(*K, t*) the amplitude of gamma range filtered white noise *ε*_*γ*_(*t*), and *n*(*t*) is the colored noise. The first two terms represent thalamic inputs (**Figure 3 A**, left). Specifically, *ε*_*γ*_(*t*) was obtained by applying a 3^rd^-order bandpass Butterworth filter of central frequency equal to 57 Hz and bandwidth equal to 10 Hz to white noise. The noise term *n*(*t*) is a z-scored colored noise, with the PSD following 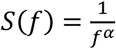, with *α* = 1.5, and an amplitude factor 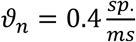. Both *ε*_γ_ (*t*) and *n*(*t*) were independently generated at every simulation. […]_+_ is a threshold-linear function, [*x*]_+ =_ *x* if *x* > 0, [*x*]_+ =_ 0 otherwise, to avoid a negative number of spikes which could arise due to the noise terms. In the first part of our work (**Figure 3** and **Figure 4**), we set the external input parameters to be time-invariant, i.e., *A*(*K*) and *S*(*K*) with a value for each K reported in Table 2 (see *Parameter selection*).

**Table 2.** Thalamic input parameters.

|  | K=0 | K=10 | K=20 | K=30 | K=50 | K=90 |
| --- | --- | --- | --- | --- | --- | --- |
| $A(K)[sp./s]$ | 40 | 30 | 20 | 0 | 0 | 0 |
| $S(K)[sp./s]$ | 500 | 500 | 500 | 500 | 600 | 700 |
| $A_0(K)[sp./s]$ | 40 | 30 | 20 | 0 | 0 | 0 |
| $\alpha(K)[sp./s]$ | 0 | 30 | 20 | 0 | 0 | 0 |
| $S_0(K)[sp./s]$ | 500 | 500 | 500 | 500 | 500 | 600 |
| $\beta(K)[sp./s]$ | 0 | 0 | 0 | 0 | 250 | 200 |

**Figure 4.**
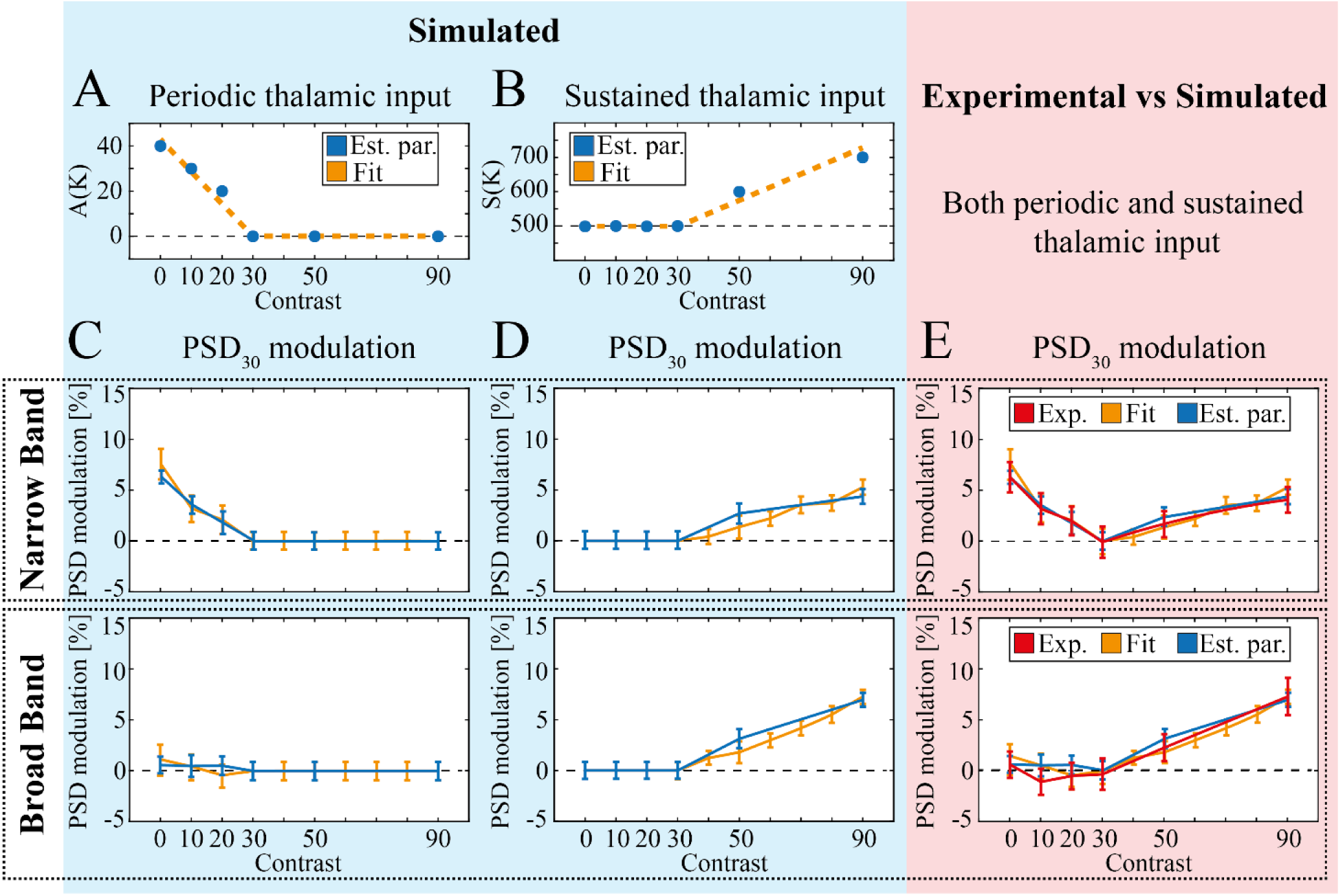
Simulation of NB and BB gamma over the whole contrast range. ***A, B***, Optimal amplitude of oscillatory (***A***) and sustained (***B***) thalamic input amplitude for each value of contrast K (blue markers) and corresponding fit (dashed orange line). ***C***, PSD_30_ modulation (see Materials and Methods) of NB (top) and BB (bottom) with fixed sustained input S=500 sp./s, and A(K) from single values shown in (***A***) (blue line) or overall fit (orange line). Error bars indicate mean ± SEM here and in the following panels. ***D***, PSD_30_ modulation (see Materials and Methods) of NB (top) and BB (bottom) with sustained input S(K) values shown in (B) (blue line) or associated fit (orange line) and A(K)=0 sp./s. ***E***, Comparison of PSD_30_ modulation in NB (top) and BB (bottom) from experimental data (red) and simulations with A(K) and S(K) values determined by local optimization (blue) or associated fit (orange).

**Figure 5.**
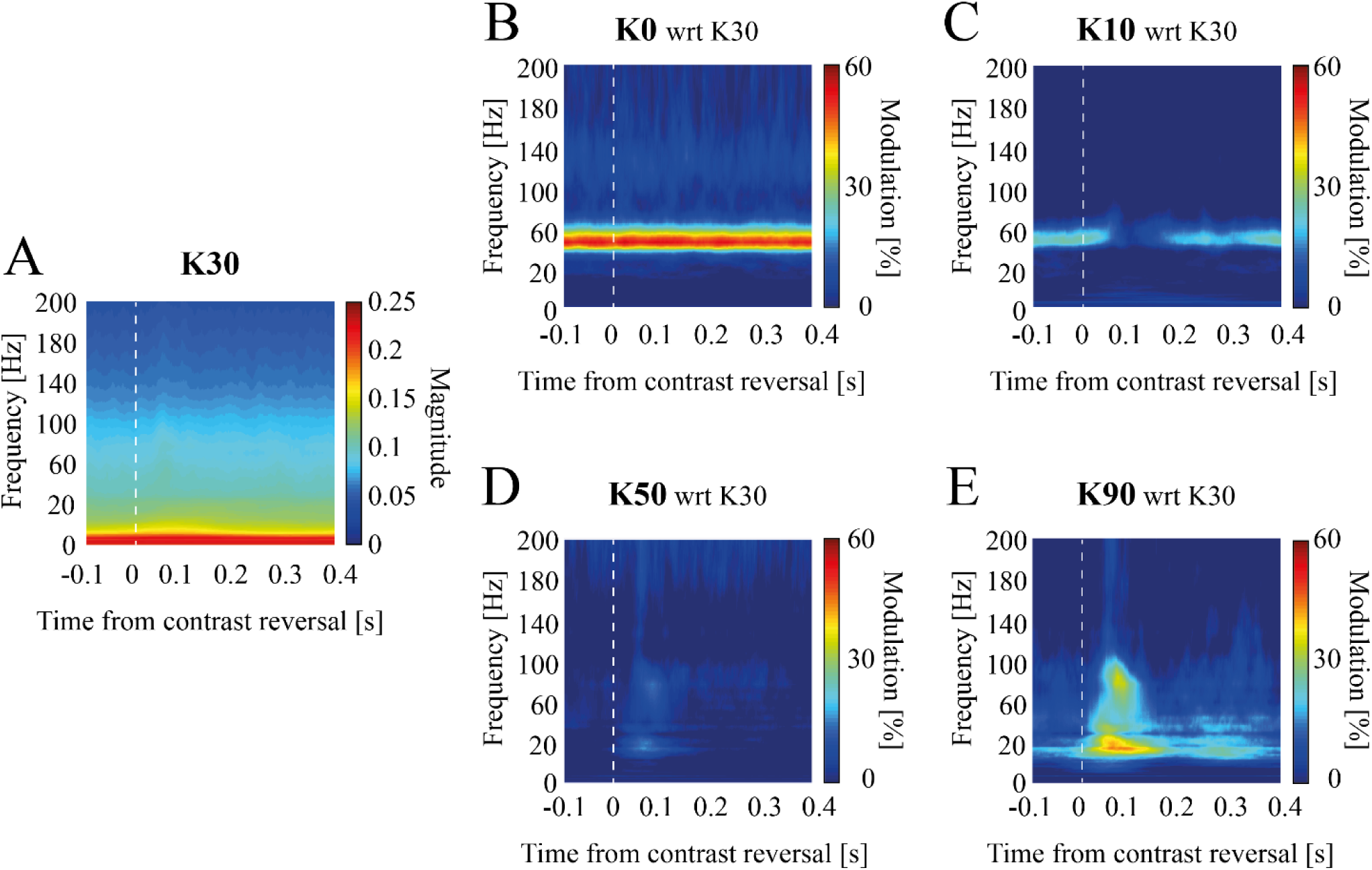
Temporal evolution of contrast-dependent spectral modulation in experimental data. Mean scalogram modulation with respect to K=30 (***A***) for experimental data at K=0 (***B***), K=10 (***C***), K=50 (***D***), and K=90 (***E***). Scalograms were averaged over trials in the interval [-100,400] ms around contrast reversals (see Materials and Methods) indicated by vertical white dashed lines.

In particular, setting 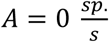 and 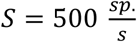 the PSD of the simulated LFPs closely matched the reference experimental spectrum for K=30 (**Figure 1 D**, middle. Reduced chi-squared between experimental and simulated LFPs PSD 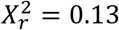). The spectral modulations of the other simulated contrast levels were consequently assessed using this median LFP PSD as a baseline, analogously to the approach adopted for the experimental data.

The values of *A*(*K*) and *S*(*K*) across K were approximated with two linear functions (**Figure 4 A-B**):

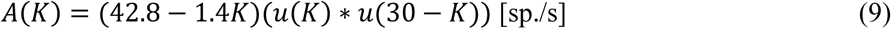

and

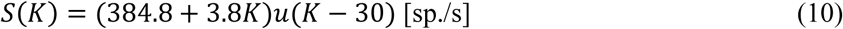

where *K* is the contrast level intended to be simulated and *u*(*K*) is the Heaviside step function: *u*(*K*) = 0 for *u*(*K*) < 0 and *u*(*K*) = 1 for *u*(*K*) ≥ 0.

In the second part of the work (**Figure 6**) we also took into account the evolution in time of the external input parameters. As for the amplitude of the gamma range filtered white noise, i.e., *A*(*K, t*), we defined it as a function of the time of grating reversal (see Visual stimuli)

**Figure 6.**
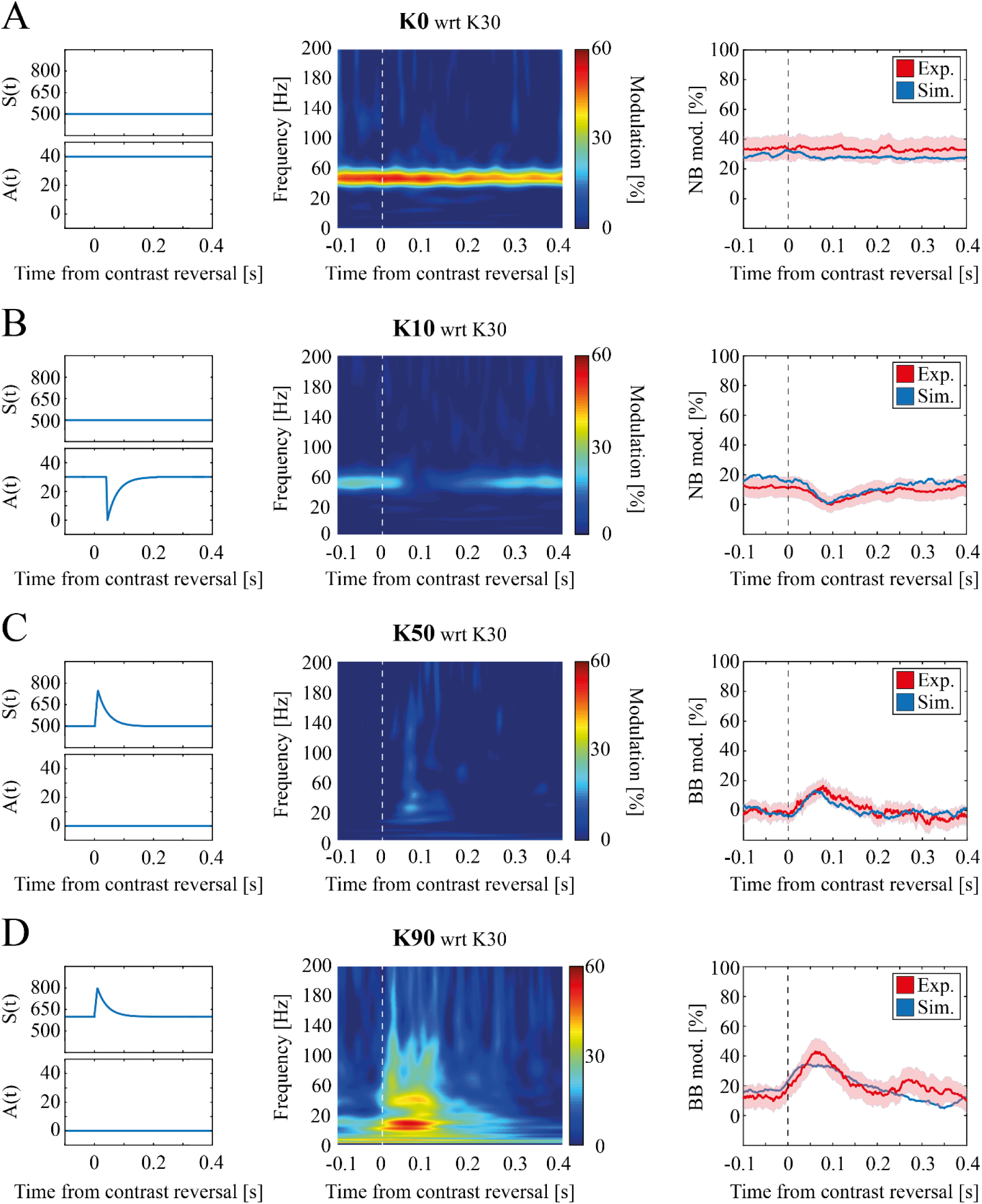
Temporal evolution of contrast-dependent modulation in simulated data. ***A***, *Left:* Time course of thalamic input model parameters for K=0. *Middle:* Mean modulation of scalogram at K=0 with respect to K=30 for simulated data. *Right:* Time evolution of narrow band modulation for experimental (red) and simulated (blue) data (shading indicates SEM). Data are averaged over trials in the interval [-100,400] ms around contrast reversals indicated by vertical white (middle) and black (right) dashed lines. ***B***, same as (A) for K=10. ***C***, *Left:* Time course of thalamic input model parameters for K=50. *Middle:* Mean modulation of scalogram at K=50 with respect to K=30 for simulated data. *Right:* Time evolution of broad band modulation for experimental (red) and simulated (blue) data (shading indicates SEM). ***D***, same as (***C***) for K=90.

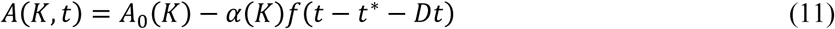

with

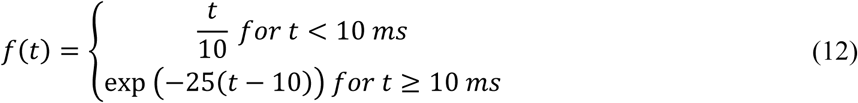

where *A*_0_(*K*) is the baseline value, *α*(*K*) is the amplitude of the reversal-driven modulation *f*(*t*), *t*^*^ is the time of grating reversal and Dt is 40 ms to mimic the latency of the narrowband observed experimentally. The finite rise time was estimated to ∼10ms (see **Figure 6 C-D**) and for the sake of simplicity was approximated with a linear growth. The decay time was defined to reproduce the quick decay observed experimentally, with the transient effects ending after ∼200 ms (see **Figure 6**). The values of *A*_0_(*K*) and *α*(*K*) are reported in Table 2 (see *Inputs parameter selection*).

Similarly, we set the sustained thalamic input *S*(*K, t*) to:

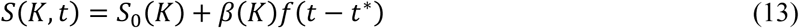

where *S*_0_(*K*) is an additive constant and *β*(*K*) is the amplitude of the same reversal-driven modulation *f*(*t*) defined in Equation (12). The values of *S*_0_(*K*) and *β*(*K*) are reported in Table 2 (see next section).

#### Input parameters selection

Starting from experimental observations (see **Figure 2 B-G**) we defined the synaptic inputs A(K) and S(K) to be complementary. Setting 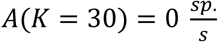 and 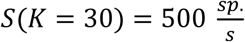 the resulting synthetic LFP spectra closely matched experimental results (see above). For all K>30 A(K) was set to be zero and for all values K<30 S(K) was set to be 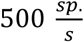. To find the optimal parameters *A*(*K*) for K = [0, 10, 20] we simulated the LFP generated by inputs with values of A(K) ranging from 0 to 100 sp./s in steps of 10 sp./ms. For each input, we estimated the spectral modulation relative to K=30 and we estimated the agreement with the experimental LFP spectral modulation as follows:

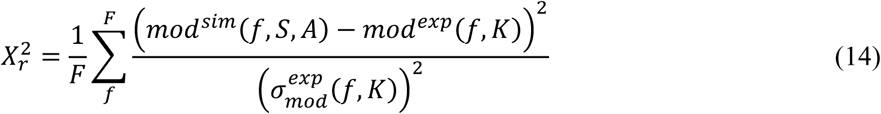

where F is the total number of frequencies, *mod*^*sim*^(*f, S, A*) is the median modulation across simulations of the LFPs when setting the external input parameters to *S* and *A* (see Equation (8)), and *mod*^*exp*^(*f, K*) and 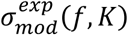 are the median and the variance modulation of the experimental LFPs at a given contrast *K* across animals and trials. For each K, the value of A(K) minimizing this measure was selected (see Table 2).

We set the optimal values of *S*(*K*) for K=[50, 90] (see Table 2) in a similar way, simulating the LFP generated by inputs in which *S*(*K*) varied from 500 to 1000 sp./s in steps of 100 sp./s.

Similarly, as we found no significant time structure in the experimental scalogram at K=30 (**Figure 5 A**), for the time-variant model we set *S*_0_(*K* = *3*0) = *5*00 *sp*.*/s, A*_0_(*K* = *3*0) = *α*(*K* = *3*0) = *β*(*K* = *3*0) = 0 *sp*.*/s*. As we have experimentally observed narrow and broad band to be complementary modulated in two different contrast ranges (**Figure 5**), for K<30 we set *β*(*K*) = 0 *sp*.*/s* and *S*_0_(*K*) = *5*00 *sp*.*/s*, whereas for K>30 we set *α*(*K*) = *A*_0_(*K*) = 0 *sp*.*/s*. Therefore, in order to find the optimal values of *S*_0_(*K*) and *β*(*K*) parameters for K=[50,90] we simulated the LFP generated by inputs with values of *S*_0_(*K*) ranging from 500 to 1000 sp./s in steps of 100 sp./ms and of *β*(*K*) ranging from 0 to 400 sp./s in eight uniform steps. For each input pair combination, we, therefore, computed the scalogram modulation relative to K=30, we extracted the BB time evolution and we estimated the agreement with the experimental LFP scalogram modulation as follows:

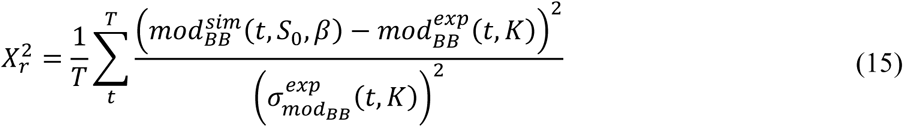

where T is the total number of time points, 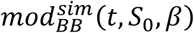 is the median modulation of the simulated time course of the broad band gamma range when setting the external input parameters to *S*_0_ and β, 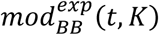 and 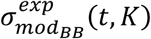 are the median and the variance modulation of the experimental time course of the broad band gamma range at a given contrast level *K*.

Similarly, for K=[0,10,20] we simulated the LFP generated by inputs with values of *A*_0_(*K*) ranging from 0 to 100 sp./s in steps of 10 sp./ms and of *α*(*K*) ranging from 0 to *A*_0_(*K*) in steps of 10 sp./s. For each input pair combination, we, therefore, computed the scalogram modulation relative to K=30, we extracted the NB time evolution and we estimated the agreement with the experimental LFP scalogram modulation as follows:

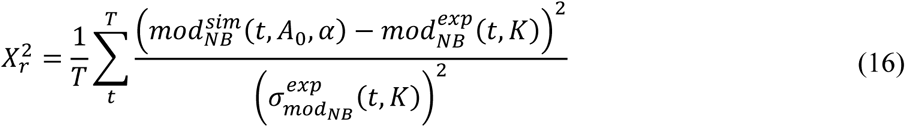

where T is the total number of time points, 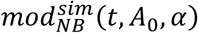 is the median modulation of the simulated time course of the narrow-band gamma range when setting the external input parameters to *A*_0_ and α, 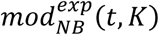 and 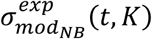 are the median and the variance modulation of the experimental time course of the narrow-band gamma range at a given contrast level *K*.

#### Simulations

Network simulations were performed using a finite difference integration scheme based on the second-order Runge Kutta algorithm (Hansel et al., 1998; Shelley and Tao, 2001; Press, 2007) with time step *Δt* = 0.0*5 ms*. To focus on stationary responses, the first 200 ms of every simulation were discarded.

Simulations with time-invariant inputs lasted 10 s and each set of inputs was presented 25 times with different noise terms *ε*_*γ*_(*t*) and *n*(*t*).

Simulations with time-dependent inputs lasted 2s: during the first second, the input was set to the baseline values of 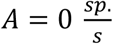 and 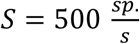; during the last second, instead, the input was determined by the level of contrast (see Table 2). Each set of input was presented 100 times with different noise terms *ε*_*γ*_(*t*) and *n*(*t*). All simulations were conducted with custom made Python scripts within the Brian 2 simulator environment (Goodman, 2008; Stimberg et al., 2019).

### Statistical analysis

Data processing and statistical analysis were carried on with custom made Matlab scripts and available third-party data analysis Toolboxes (e.g., for computing mutual information). We employed both custom scripts and built-in data analysis and statistical functions. For each statistical comparison throughout the text, we reported the statistical test and their p-values. P-values lower than 0.05 were considered significant. All results will be reported as median ± SEM unless otherwise stated

## Results

### Broad and narrow gamma bands in mice V1 display distinct sensitivity to contrast

We analyzed the spectral proprieties of the LFP recorded from V1 layer IV of awake mice presented with alternating gratings with different levels of contrast (see Materials and Methods and **Figure 1 A**). Coherently with previous results (Saleem et al., 2017), we found that spectral modulation of the LFP with such stimuli was not uniform over the whole gamma band [20-95] Hz, but was characterized by the presence of a narrow (NB, [45-65] Hz) and a broad band (BB, [20-45] Hz and [65-95] Hz) with distinct modulation proprieties. Both bands increased their power at stimulus onset (i.e., screen ON with simultaneous luminance and contrast variation; **Figure 1 B**, Wilcoxon’s matched pairs signed rank test p<<0.001 for both NB (top) and BB (bottom). The subsequent response (i.e., to contrast reversal) of NB was maximal for low contrasts (K=0, see **Figure 1 C**, top) while for BB was maximal at high contrasts (K=90, see **Figure 1 C**, bottom). As a consequence, the two bands displayed opposite spectral modulation compared to intermediate contrasts (K=30, see **Figure 1 D**).

We quantified the response over time of narrow band and high ([65-95] Hz) and low ([20-45] Hz) broad band (**Figure 2 A**). NB was significantly modulated by visual contrast throughout the whole inter-contrast-reversal interval, i.e., 500 ms (**Figure 2 B** top, Kruskal-Wallis One-Way ANOVA (K-W ANOVA1): p<<0.01). Specifically, NB at very low contrast (K=0) had a significantly higher power than at intermediate contrast (K=30) (K=0: [-33.62±0.52] dB/Hz vs K=30: [-36.33±0.49] dB/Hz, median ± SEM, post-hoc Dunn’s test, p<<0.01, **Figure 2 B** top) while no significant modulation in power spectral density was found between contrasts from K=30 to K=90 (K=30: [-36.33±0.49] dB/Hz vs K=90: [-35.78±0.73] dB/Hz, post-hoc Dunn’s test, p>0.5, **Figure 2 B** top). We repeated the analysis separately for the intervals [0-200] ms and [200-500] ms following contrast reversal to discriminate between early responses due to reversal and late responses due to the presentation of static contrast. NB PSD modulation was similar across these two time windows (**Figure** 2 **C**, left, K-W ANOVA1: p<<0.01; **Figure 2 C**, right, K-W ANOVA1: p<<0.01).

Low range BB ([20-45] Hz) was significantly modulated by contrast throughout the whole inter-contrast-reversal interval (**Figure 2 B** middle, K-W ANOVA1: p=0.02). Unlike NB, low BB power was uniform for levels of contrast K≤30 (post-hoc Dunn’s test, p>0.5), while increased significantly at high contrast (K≥30) (K=90: [-28.53±0.74] dB/Hz vs K=30: [-31.26±0.49] dB/Hz, post-hoc Dunn’s test, p=0.04, **Figure 2 B** middle). Low BB early response was significantly modulated by contrast level (**Figure 2 D** left, K-W ANOVA1: p=0.02), with the highest contrast (K=90) displaying significantly higher power than at intermediate contrast level (K=90: [-26.90±0.80] dB/Hz vs K=30: [-30.84±0.48] dB/Hz, post-hoc Dunn’s test, p=0.03, **Figure 2 D** left). The low BB late response, instead, was not significantly modulated by contrast (**Figure 2 D** right, K-W ANOVA1 p=0.06).

High range BB ([65-95] Hz) was not significantly modulated by contrast throughout the whole inter-contrast-reversal interval (**Figure 2 B** bottom, K-W ANOVA1: p=0.46). This range was only significantly modulated by contrast reversal (**Figure 2 E** left, for the early response, K-W ANOVA1: p=0.04. K=90: [-35.68±0.71] dB/Hz vs K=30: [-38.59±0.46] dB/Hz, post hoc Dunn’s test, p=0.04. **Figure 2 E** right, for the late response, K-W ANOVA1: p=0.30).

Overall these results show that: (i) high and low broad band have similar behavior and therefore in the following will be considered as one broad band (BB, as in (Saleem et al., 2017)); (ii) narrow and broad band power are respectively anticorrelated and correlated with contrast; (iii) narrow (broad) band modulation occurs primarily at late (early) stage following contrast reversal.

The complementary sensitivity to visual contrast of narrow and broad band was furtherly investigated by examining the mutual information carried by the spectral modulation with respect to K=30 of the two gamma bands. We found NB to carry significant information in the low contrast range below K=30 both throughout the whole inter-contrast-reversal interval (**Figure 2 F**, top, information above significance threshold was achieved considering every contrast level ‘All K’ and the lower contrast range ‘K<30’) and from 200 to 500 ms after contrast reversal (**Figure 2 F**, bottom, information above significance threshold was achieved considering every contrast level and both the lower and the upper range of contrast levels), while information was reduced for K>30. However, within 200 ms after contrast reversal (**Figure 2 F**, middle), the NB modulation did not exceed significance threshold in any contrast level range. Conversely, we found BB to carry greater information in the higher contrast range above K=30 throughout the whole inter contrast reversal interval (**Figure 2 G**, top, information above significance threshold was achieved considering the upper range of contrasts K>30) and within 200 ms following contrast reversal (**Figure 2 G**, middle, information above significance threshold was achieved considering every contrast level and both the lower and the upper range of contrast levels), while no significant information was detected when considering a time window spanning from 200 ms to 500 ms after contrast reversal (**Figure 2 G**, bottom).

The spectral modulations of the two bands displayed a synergistic contribution in the encoding of visual contrasts in all of the temporal windows considered (synergy=0.18 for [0-200] ms; synergy=0.39 for [200-500] ms; synergy=0.20 for [0-500] ms).

Altogether these results show NB and BB to display an opposite behavior, with the former encoding increase in contrast from K=0 to K=30 in a progressively decreasing power, and the latter encoding increase in contrast from K=30 to K=90 in a progressively increasing power. To highlight this result, in the following we will employ the reference contrast value K=30 (whose PSD is reported in **Figure 1 D** middle) to discriminate between these two different modulations, as this contrast value induces minimal power for both NB and BB.

### Spiking neuronal network captures distinct functional gamma bands

To model the modulation with contrast of these functional gamma bands in V1, we built on previous works of our group accounting for BB dynamics (Mazzoni et al., 2011) to include periodic thalamic inputs associated with NB (Saleem et al., 2017) (see Materials and Methods). Briefly, the simulated input was given by the superimposition of two components: a sustained one with an average intensity depending on contrast (S(K)), and an oscillatory one with a contrast dependent amplitude (A(K)) (see **Figure 3 A**, left and Equation (8)). The model reproduced quantitatively (modulation at 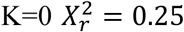; modulation at 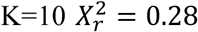) both the location and the shape of the NB frequency peak for different levels of contrast (**Figure 3 B**). This was achieved by varying A(K) while S(K) was fixed (see Table 2). For the same set of network parameters (see Table 1), the model quantitatively (modulation at K=50 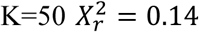; modulation at K=90 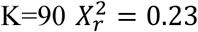) reproduced BB peak shape and BB power increase from reference to high contrast (**Figure 3 C**). This was achieved by varying S(K) while A(K) was fixed (see Table 2). This suggests that the functional modulation of these two gamma-bands can be accounted for by a proper selection of the two parameters defining the thalamic inputs, with S(K) determining the intensity of the BB and A(K) determining the intensity of the NB.

### Complementary contribution of oscillatory and sustained thalamic input to LFP gamma activity

We explored then the possibility of capturing in the model the whole range of the contrast-dependent gamma modulation by defining two complementary ranges of contrast sensitivity for the oscillatory input A(K) and sustained input S(K). We set the oscillatory input A(K) to zero in the range K≥30, while in the K<30 range we selected for each K the value of A maximizing the overall similarity between simulated and experimental spectral modulation (see Materials and Methods), obtaining a monotonous negative trend (see Table 2) that could be described by a linear fit (see **Figure 4 A** and Equation (9)). When coupled with a fixed value of S(K) this A(K) led to a decrease of the narrow band with K (**Figure 4 C** top, Kruskal-Wallis ANOVA p>0.9 for K≥30, Kruskal-Wallis ANOVA p=0.08 for K≤20, Kruskal-Wallis ANOVA p<0.01 for K≤10) and negligible fluctuations in the broad band (**Figure 4 C** bottom, Kruskal-Wallis ANOVA p>0.9).

Conversely, we set the sustained input S(K) to have a fixed baseline value S(K)=500 sp./s up to K=30 while in the contrast range K>30 for each K we selected the value of S maximizing the overall similarity between the simulated and experimental spectral modulation, obtaining a monotonous positive trend that was described by a linear fit (**Figure 4 B** and Equation (10)). This led to an increase of both NB (**Figure 4 D** top, Kruskal-Wallis ANOVA p=0.06 for K≤70 and Kruskal-Wallis ANOVA p<<0.001 for K≤90) and BB (**Figure 4 D** bottom, Kruskal-Wallis ANOVA p=0.09 for K≤60 and Kruskal-Wallis ANOVA p<<0.001 for K≤90) for K increasing in the high contrast range (see also **Figure 1 C** bottom and Discussion).

We expect A(K) and S(K) to be present at each time for any level of input in real conditions. To compare experimental and simulation results over the whole range of contrasts we then injected in the network for each value of K the input resulting from the superimposition of both A(K) and S(K) values as defined above. We found that the network reproduced the modulation of both the NB and the BB over the whole set of contrasts (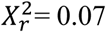 and 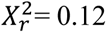, **Figure 4 E** top and bottom respectively). Similar results were achieved replacing the values of A(K) and S(K) optimized for each value of K with two linear functions (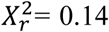 and 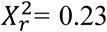, **Figure 4 E**, top and bottom respectively). These results show (i) that the dynamics of gamma bands in mouse V1 can be reproduced by a simple network architecture with a proper design of thalamic inputs, (ii) that very good agreement with experimental data is achieved by modeling thalamic input as the sum of an oscillatory component, sensitive to low levels of contrast, and a sustained component, sensitive, instead, to high levels of contrast. A broader exploration of input parameters revealed that in general NB is sensitive to the increase in the oscillatory input A(K) and, to a lesser extent, to the increase in the sustained input S(K) (Two-way ANOVA showed p<<0.001 for parameter A, p<<0.001 for parameter S and p=0.3 for their interaction), while BB is only sensitive to the sustained input S(K) (Two-way ANOVA showed p=0.33 for A, p<<0.001 for S and p=0.32 for their interaction).

### Narrow and broad band gamma dynamics at contrast reversal

As we observed differences in early and late response, we focused on the temporal evolution of NB and BB (see Materials and Methods). As expected, narrow band modulation at K=0 showed a sustained activity throughout the whole stimulation with no specific response associated with contrast reversal (**Figure 5 B**). For K=10, NB power decreased after the grating contrast reversal (minimum NB modulation: [-4.5±2.3]% reached at [107±5] ms, **Figure 5 C**). This decrease in NB power lasted for about 200 ms (K=10 offset: [198±10] ms). Broad band gamma power, instead, expressed a rapid transient increase immediately after high contrast reversal (maximum BB modulation at K=50 [17±4] % reached at [73±8] ms, **Figure 5 D**. Maximum BB modulation at K=90 [46±9] % reached at [76±9] ms, **Figure 5 E**). After this transient enhancement, BB power returned to the pre-contrast reversal baseline level within 200 ms (as previously observed for the NB; K=50 offset: [177±2] ms; K=90 offset: [209±1] ms).

Of note, NB and BB displayed different temporal dynamics relative to contrast reversal, with the former having a much longer delay (NB onset: [42±5] ms for K=10; BB onset: [0-16] ms, [25^th^-75^th^] percentile, for K=50 and K=90; Wilcoxon’s rank-sum test p<0.05).

### Time-dependent thalamic input model accounts for gamma band temporal evolution

Modeling results shown in **Figure 3-4** reproduce the average response of the network over time. To reproduce the experimental time-frequency features of V1 LFP we re-defined the simulated thalamic input to be also a function of time (see Equation (12)) by taking into account a reversal-driven modulation at contrast reversal. The function was defined in such a way to be zeroed after 200 ms (as observed experimentally in the LFP scalograms, **Figure 6** last column). We then re-determined A(K,t) and S(K,t) to optimally fit experimental data also over time rather than only averaged over the whole interval (see Materials and methods). This allowed capturing the differences in early and late thalamic stimulation leading to the distinction of early and late cortical responses.

By introducing this temporal dynamic in the thalamic input we were able to reproduce the experimental LFPs at contrast reversal, both the NB time invariance at K=0 (**Figure 6 A** middle, and 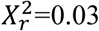 in **Figure 6 A** right), the NB power decrease at K=10 (**Figure 6 B** middle, and 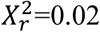 in **Figure 6 B** right. 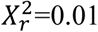 for K=20, data not shown) and the BB power increase for high contrast (K=50: **Figure 6 C** middle, and 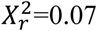 in **Figure 6 C** right. K=90: **Figure 6 D** middle, and 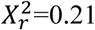 in **Figure 6 D** right).

## Discussion

We found that, in V1 of awake mice, narrow and broad gamma band displayed an opposite modulation with contrast and were sensitive to complementary contrast ranges. Starting from these experimental results we developed a simple leaky integrate-and-fire spiking network model of mice V1, which reproduced quantitatively the observed dynamics of contrast-driven gamma modulations, providing that the thalamic input is composed by the superimposition of a sustained and an oscillatory component.

### Contrast and luminance sensitivity

This work focused on the V1 processing of visual contrast. The narrow band (NB) was found to be sensitive to variations among low levels of contrast. However, we observed a small and non-informative increase of the NB power for high levels of contrast. Our model suggests that this increase might be due to the overall increase in the whole gamma 20-90 Hz range, which includes the NB and it might be due to cortical resonance rather than to an increase in the oscillatory LGN activity, coherently with (Saleem et al., 2017). The positive peak of BB associated with reversal of high contrast gratings (**Figure 6 C-D**) is coherent with previous results, suggesting that BB is sensitive to both spatial and temporal contrast (Barbieri et al., 2014). On the other hand, the NB vanishing is associated with reversal of low contrast bars (**Figure 6 B**), suggesting that NB power is negatively correlated to both spatial and temporal contrast.

One of the main limitations of this study is that we did not explicitly tackle the luminance sensitivity of the two bands. Indeed, Saleem and co-authors found that LGN gamma narrow band was also modulated by light intensity, with a higher luminance being associated with a narrow band with larger power and higher peak frequency (Saleem et al., 2017). We performed experiments with fixed luminance, hence we could not observe the consequences of the luminance modulation in V1. Our model predicts that the cortical NB oscillation is proportional to the gamma oscillatory component of the LGN input. Hence, we would expect the light-induced increase in LGN gamma activity to drive proportional increases in cortical NB, coherently with (Saleem et al., 2017). Indeed, we observed that stimulation onset was associated with an increase in both NB and BB (**Figure** 1 B), coherently with what was reported by (Saleem et al., 2017).

### Laminar gamma in rodents

Another limitation of this work is that we focused only on the analysis of the thalamocortical recipient layer IV of V1 (see Methods). Gamma activity is known to spread non homogeneously across layers both when it is stimulus-unrelated (Welle and Contreras, 2016; Senzai et al., 2019) and stimulus-related (Xing et al., 2012; Bastos et al., 2014; van Kerkoerle et al., 2014; Saleem et al., 2017). We decided here to focus on layer IV, as this layer receives direct thalamic inputs and displays early and strong gamma oscillations when presented with visual contrast stimuli (Saleem et al., 2017). Future works will investigate specific properties of propagation of the two bands observed here, analyzing the activity of different recording sites in laminar electrodes and exploiting the knowledge of inter-laminar interactions of advanced models of mice V1 (see next subsection).

### Models of mouse visual cortex

Recent models of mouse visual cortex ((Billeh et al. 2020), but see also (Troyer et al., 1998; Arkhipov et al., 2018)), succeeded in reproducing firing rate distribution across cortical layers in response to arbitrary visual inputs with a multi-layer network composed by multi-compartment generalized LIF neurons. In these models LGN inputs are modeled as a series of spatio-temporal filters of the presented image and lack the gamma band activity observed in LGN (Saleem et al., 2017; McAfee et al., 2018). Gamma modulation due to visual stimuli is indeed present in the model only with a sensitivity similar to the experimentally observed broad band, coherently with our hypothesis that the anti-correlation between V1 narrow band power and contrast might originate from periodicity in the LGN input. Combining our proposed model of LGN inputs with multi-layer models as (Billeh et al. 2020) could unravel the specific inter-layer dynamics of the narrow band.

A recent interesting modeling study reproduced the effects of LGN input modulation on V1 gamma responses to visual stimuli (Zachariou et al., 2019) with a single-layer network of conductance-based spiking neurons. However, the model aimed at reproducing the saturation of gamma power at mid-level of visual contrast that has been observed in human and non-human primates (Hadjipapas et al., 2015) and was not found in our data nor similar murine data (Saleem et al., 2017; McAfee et al., 2018).

### Narrow and broad gamma band in primates

Encoding of contrast in the V1 gamma band is known to be present in primates (Henrie and Shapley, 2005). A comparison of the role of narrow and broad gamma band between rodents and primates studies is hampered by the fact that nomenclature is different in the two cases. In primates studies narrow and broad gamma band are defined as low vs high gamma ranges: [30 80] vs [80 200] Hz (Ray and Maunsell, 2010), [20-60] vs [70 150+] Hz (Bartoli et al., 2019), or even as overlapping [30-80] vs [30-200] Hz (Hermes et al., 2019).

Many studies have observed that a sudden increase in contrast leads to a transient broad band activation followed by a sustained narrow band activity (Swettenham et al., 2009; Ray and Maunsell, 2010; Xing et al., 2012b, 2012a; Murty et al., 2018; Shirhatti and Ray, 2018; Peter et al., 2019). Unlike mice, however, the power and the peak frequency of the sustained gamma narrow band were found to be positively modulated by visual contrasts and invariant to luminance variation (Peter et al., 2019).

The V1 spiking neurons network we adopted here is not significantly different from the one we previously proved to reproduce contrast modulations in primates V1 (Mazzoni et al., 2008, 2010, 2011), except for the different thalamic inputs. This implies that we would expect to observe NB as defined in this manuscript also in primates if NB oscillations were present in primates’ LGN. However, while the sustained narrow gamma oscillation has been reported in pre cortical structures such as LGN or even the retina for rodents and cats ((Neuenschwander and Singer, 1996; Storchi et al., 2017), simultaneous recording from primate V1 and LGN found no narrow band in V1 and no gamma oscillation in LGN (Bastos et al., 2014).

### Perspectives

Mice are becoming the favorite animal model to study the circuit changes involved in several neurological disorders. This is due to the availability of sensitive imaging techniques and opto- and chemo-genetic approaches which allow identifying the cellular underpinnings of the disease (Götz et al., 2018; Fagiolini et al., 2020). Mice are also becoming a standard model for the visual cortex (Busse et al., 2011; Carandini and Churchland, 2013; Haider et al., 2013; Saleem et al., 2017). In this context, the monitoring of visual responses represents a promising biomarker for preclinical and clinical studies on neurodevelopmental disorders (LeBlanc et al., 2015; Lupori et al., 2019). Thus, an effective in silico model reproducing cardinal functions of the mouse visual cortex could lay the ground for a better understanding of the pathogenetic mechanisms underlying functional impairment in neurological diseases. In future works, we aim at investigating disorders involving the visual cortex with an interplay of experimental and modeling studies starting from the results presented here.

## Acknowledgments

This work was supported by the Italian Ministry of Research (MIUR, PRIN2017, PROTECTION, project 20178L7WRS).

